# Two routes to value-based decisions in Parkinson’s disease: differentiating incremental reinforcement learning from episodic memory

**DOI:** 10.1101/2024.05.03.592414

**Authors:** Leila Montaser-Kouhsari, Jonathan Nicholas, Raphael T. Gerraty, Daphna Shohamy

## Abstract

Patients with Parkinson’s disease are impaired at incremental reward-based learning. It is typically assumed that this impairment reflects a loss of striatal dopamine. However, many open questions remain about the nature of reward-based learning deficits in Parkinson’s. Recent studies have found that a combination of different cognitive and computational strategies contribute even to simple reward-based learning tasks, suggesting a possible role for episodic memory. These findings raise critical questions about how incremental learning and episodic memory interact to support learning from past experience and what their relative contributions are to impaired decision-making in Parkinson’s disease. Here we addressed these questions by asking patients with Parkinson’s disease (n=26) both on and off their dopamine replacement medication and age- and education-matched healthy controls (n=26) to complete a task designed to isolate the contributions of incremental learning and episodic memory to reward-based learning and decision-making. We found that Parkinson’s patients performed as well as healthy controls when using episodic memory, but were impaired at incremental reward-based learning. Dopamine replacement medication remediated this deficit while enhancing subsequent episodic memory for the value of motivationally relevant stimuli. These results demonstrate that Parkinson’s patients are impaired at learning about reward from trial-and-error when episodic memory is properly controlled for, and that learning based on the value of single experiences remains intact in patients with Parkinson’s disease.

## Introduction

The striatum and its dopaminergic inputs are thought to support a specialized circuit for incremental reward-based learning.^1,2^ It is commonly thought that this is the cause of learning deficits in patients with Parkinson’s disease (PD). Indeed, PD is characterized by a loss of striatal dopamine and PD patients have been shown to be impaired at incremental reward learning.^3–6^ However, recent findings call this simple explanation for learning deficits in PD into question, revealing that incremental learning involves a multitude of cognitive and computational strategies.^7,8^ Thus, open questions remain about the nature of learning deficits in PD and their attribution to different cognitive and computational systems.

In particular, recent work has demonstrated that tasks typically thought to measure incremental learning may instead depend on memory for one-shot events, referred to as “episodic memory”.^9– 11^ This may be especially relevant for understanding learning deficits in PD, because some studies have shown that PD patients present with deficits in episodic memory.^12–18^ This raises the question: what are the relative contributions of episodic memory and incremental learning to impaired reward-based learning in PD? Characterizing the relationship between these systems remains an important goal for understanding the exact nature of PD patients’ reward learning deficits, and for understanding the role of striatal dopamine in learning from past experience more broadly.

Incremental learning and episodic memory are classically understood as two separate learning systems with distinct neural substrates. Incremental learning depends on reward prediction errors encoded by phasic dopamine, which aid in the formation of stimulus-action associations in the striatum.^2^ Computational modeling suggests that this type of learning depends on summarizing past experiences with a running average, providing a mechanism for evaluating actions without needing to maintain individual memory traces.^1,19,20^ In contrast, episodic memory supports the encoding and retrieval of single experiences and largely depends upon independent neural circuitry, including the hippocampus.^9–11,21–27^

Understanding how these systems each contribute to performance on reward-based learning and decision tasks has been challenging, in part because typical experiments only measure one type of behavior. In particular, most tasks require participants to choose between a set of repeated and often highly familiar options. This approach is well-suited for measuring incremental learning but offers no way to test whether individual events are referenced during choice. Recent experimental approaches have offered a more sophisticated way to measure both forms of learning by associating unique cues with each option, allowing for direct measures of the role of episodic memory in reward-based learning.^10,11,24,25,28^ Importantly, such studies have revealed that tasks that may appear at first glance to depend only on incremental learning instead also depend substantially on episodic memory.

Could it be then that episodic memory impairments may also contribute to learning deficits in PD? It is typically assumed that patients with PD are primarily impaired at incremental learning from feedback, due to dysfunction in striatal reward signals. However, recent results have complicated this interpretation in a number of ways. First, some studies have found that incremental reward learning is sometimes preserved in PD patients.^31,32^ Second, studies in healthy adults have reported a relationship between BOLD activity in the striatum and episodic retrieval.^18,27,37–42^ These findings suggest that patients with PD may have episodic memory impairment. Indeed, some studies in PD have shown that patients have deficits in episodic recall,^12–17^ although the evidence is mixed, with others finding little-to-no issues.^33–36^ Together, these findings suggest that there could be some shared dependence of episodic memory and incremental learning on striatal dopamine. Yet because most tasks in PD patients measure only episodic memory or incremental learning alone, it is currently unclear whether either system, or even both, is responsible for these deficits. The involvement of episodic memory in reward-based learning, and the possibility of its disruption, makes disentangling its unique contributions from those of incremental learning critical to our understanding of learning deficits in PD.

Here, our goal was to address this gap by determining the extent to which PD patients use either incremental learning or episodic memory to guide reward-based learning. We used an experimental task that was designed to provide clear behavioral measures of participants’ use of either of these strategies (**Figure 1**).^25^ PD patients (N=26) completed this task while on- or off-dopamine replacement medication. Their performance was compared to a group of matched healthy controls. Following the task, both groups were given a subsequent memory test. This design allowed us to determine whether PD patients were impaired at using either incremental learning or episodic memory to guide reward-based learning, and whether dopamine replacement remediated any deficits.

**Figure 1.**
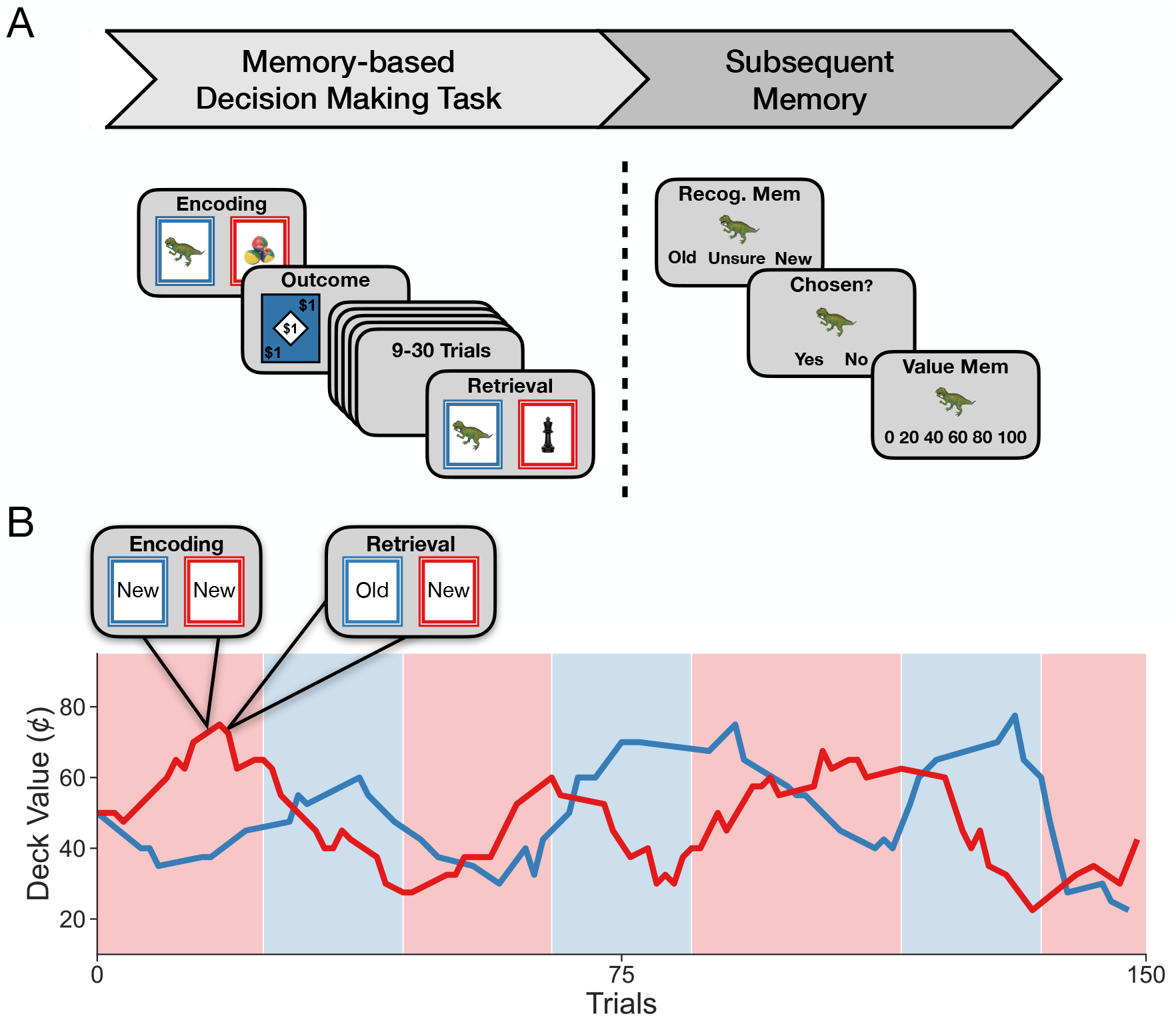
Design of the experiment. **A)** Participants completed two tasks in succession. The first was the *memory-based decision-making task*. Participants chose between two colored cards and received an outcome following each choice. One color was worth more on average at any given timepoint and this mapping changed periodically. Each card also displayed a trial-unique object, and cards that were chosen could appear a second time in the task after 9-30 trials. If a card re-appeared, it was worth the same amount, which allowed participants to use episodic memory for individual cards in addition to learning deck value from feedback. Outcomes ranged from $0 to $1 in increments of 20¢. Participants then completed a *subsequent memory* task for objects that may have been seen in the memory-based decision-making task. Participants had to indicate whether they recognized an object and, if they did, whether they chose that object. If they responded that they had chosen the object they were then asked if they remembered the value of that object. **B)** Participants completed 150 trials, with the higher value deck alternating periodically (every 10-24 trials), as indicated by the shaded background. Lines indicate an example of tracking the value of each deck according to outcomes received for a single participant.

We aimed to test three hypotheses. Patients could be impaired at either:

i. Incremental learning alone. This would be expected if a lack of striatal dopamine results in dysfunctional reward prediction error with no effect on episodic memory.
ii. Episodic memory alone. This would be expected if prior tasks have mischaracterized episodic retrieval deficits as incremental learning deficits.
iii. Both systems simultaneously. This would be expected if there is shared dependence on striatal dopamine by both systems.

Finally, we also asked whether dopamine replacement had any impact on episodic memory formation by examining patients’ memory for individual events following reward learning.

## Materials and Methods

### Participants: PD patients and healthy controls

Twenty-six patients with PD (aged 50-80; 34% women) were recruited from the Columbia University Neurological Institute, the Columbia University Recruit Me subject pool, or the Michael J Fox Foundation Trial Finder website. Twenty-six healthy controls (HC; aged 50-80; 53% women) were also recruited from the local community. Patients were age and education matched with healthy controls (there were no significant differences between groups on these variables) and had no history of other neurological diseases besides PD and no major active psychiatric disorders. Patients were all in the mild-to-moderate disease stage: mean UPDRS off 27.43 ± 9.78, as examined by a movement disorders neurologist; disease duration 2-14 years (**Table 1**). Patients were being treated with carbidopa/levodopa for at least one year prior to testing and were responsive to their dopaminergic medication. While our sample included more men than women in the PD group compared to healthy controls, this was unsurprising given the higher prevalence of PD in men. Informed consent was obtained with approval from the Columbia University Institutional Review Board.

**Table 1.** Demographic and clinical characteristics of participants.

|  | Healthy controls | Parkinson's patients | P-value (FDR-corrected) |
| --- | --- | --- | --- |
| Age (years) | 65.84 | 66.11 | 1.0 |
| Sex (% female) | 53 | 34 | 1.0 |
| Education (years) | 16.38 | 17.41 | 0.513 |
| MoCA | 28.23 | 29 | 0.391 |
| Geriatric Depression Scale | 3.39 | 6.98 | 0.117 |
| Starkstein Apathy Scale | 5.73 | 9.65 | 0.029 |
| Digit Span forward<sup>a</sup> | 11.80 | 12.69 | 0.537 |
| Digit Span backward | 7.88 | 7.88 | 1.0 |
| Digit Span forward (off) | -- | 12.69 | 0.537 |
| Digit Span backward (off) | -- | 7.84 | 1.0 |
| UPDRS off | -- | 27.03 | -- |
| UPDRS on<sup>b</sup> | -- | 18.46 | -- |
| Disease duration (years) | -- | 6.657 | -- |

Table shows mean values. MoCA = Montreal Cognitive Assessment; UPDRS = Unified Parkinson's Disease Rating Scale—Part III.
<sup>a</sup>First two digit span rows indicate scores for Parkinson's patients on medication
<sup>b</sup>UPDRS on was significantly lower than UPDRS off ( $p < 0.001$ )

### Procedure

Before the experimental task, all participants completed a neuropsychological battery lasting approximately 30 minutes comprised of the Montreal Cognitive Assessment (MoCA), geriatric depression scale (GDS), digit forward and backward span, and the Starkstein apathy scale. This battery was selected based on the current understanding of PD non-motor symptoms, which can include depression, executive function, and apathy. Participants with mild cognitive impairment or dementia as indicated by a MOCA score less than 26 were not included in the study.

PD patients performed the experiments in 2 sessions that were between 1-7 days apart, once on-medication, once off-medication. The order of the on and off sessions was counterbalanced across patients. For the on-medication session, patients took their medication 1–1.5 hours before behavioral testing. For the off-medication session, patients withdrew from their medication overnight (at least 12 hours for levodopa/carbidopa and at least 24 hours for dopamine agonists). All participants performed several tasks in each session, counterbalanced in order.

### Tasks

The primary task was developed by our lab to measure the relative contributions of incremental learning and episodic memory to value-based decisions (**Figure 1**).^24,25^ On each trial, participants were presented with two decks of cards (red and blue) and had 2 seconds to choose between them (using either the “j” or “k” keys). They were then shown the outcome of their choice (i.e., the selected card’s value) for 1.5 seconds, followed by a 1 second intertrial interval. Each card’s value was between $0 and $1 with $0.20 increments. Deck position on the screen (left or right) was counterbalanced, and participants completed 150 trials in total.

Participants were made aware that one deck color had a higher expected value at any given time and was therefore the “lucky” deck (i.e., *V*_*lucky*_ *=* $0.63, *V*_*unlucky*_ *=* $0.37). Importantly, the currently lucky deck reversed frequently throughout the task (every 16-24 trials), requiring participants to use each deck’s recent reward history to determine the identity of the lucky deck. Each card within a deck featured an image of a trial-unique object which varied on each trial, and a subset of chosen images (60%) appeared once more throughout the task. This allowed participants to use episodic memory for an individual image: Participants were told that if they encountered a specific image on a card a second time, it would be worth the same as when the card with that image was first chosen, regardless of whether its deck color was currently lucky or not. The repeated cards appeared 9-30 trials after the first presentation and were worth the same amount as the first time they were chosen. Objects were sampled using a procedure designed to prevent each deck’s expected value from becoming skewed by choice to minimize the correlation between the expected value of previously seen cards and deck expected value and ensure that choosing a previously selected card remained close to 50¢ (the sampling procedure is described in previous publications^24,25^).

Following completion of the decision-making task, participants underwent a memory test for a subset of the trial-unique objects. The memory test followed a 5-10 minute delay during which instructions and practice trials were completed. For each trial of the memory test, an object was first displayed on the screen and participants were asked whether they had previously seen the object and were given five response options: *Definitely New, Probably New, Don’t Know, Probably Old, Definitely Old*. If the participant reported that they had not seen the object before or did not know, they moved on to the next trial. If they reported that they had seen the object before, they were then asked if they had chosen the object or not. Lastly, if they responded that they had chosen the object, they were asked what the value of that object was (with options spanning each of the six possible object values between $0-1). Participants completed up to 99 such trials in total.

### Analyses

To quantify estimates of incrementally constructed value in the memory-based decision-making task, we modeled learning about deck value using a modified Rescorla-Wagner^20^ model. This model assumes that each participant’s stored expected value, *Q*, for each deck, *d*, is first updated on each trial, *t*, via a prediction error, *δ*_t_, representing the difference between the expected and received outcome (*R*_*t*_), weighted by a learning rate α:

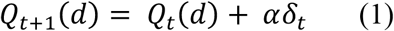

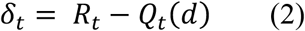

And is not updated if a different option is chosen:

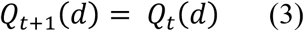

Outcomes were centered to lie between -0.5 and 0.5. To quantify the influence of episodic memory on decision-making, we included the object value for trials featuring previously seen objects trials (*OldValue*; coded to range from 0.5 if the value was $1 on the red deck or $0 on the blue deck to –0.5 if the value was $0 on the red deck and $1 on the blue deck) in a sigmoid choice function alongside deck expected value. We additionally accounted for other potential influences on choice by including 1) “familiarity” (Fam) as a binary variable indicating whether a choice included (coded as 0.5) or did not include (coded as -0.5) a previously seen object on one of the two decks, 2) “perseveration” (*Perse*) as a binary variable used to capture any bias toward repeating a previous choice independent of reward outcome (coded as 0.5 if a choice was repeated and -0.5 if it was not), and 3) an intercept representing a baseline bias toward either color of deck. A vector of five inverse temperature parameters was included to model these effects for each participant: a deck color bias *β*_0_, an effect of expected deck value *β*_1_, an effect of episodic object value *β*_2_, an effect of familiarity bias *β*_3_, and an effect of perseveration bias *β*_4_. These inverse temperatures were used to model the probability of choosing the red deck:

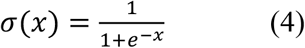

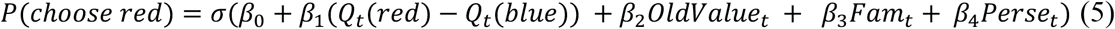

Hierarchical Bayesian inference was used to fit parameters for individual participants within each group in order to pool uncertainty across the entire sample and aid in identifiability^45^. Learning rate and inverse temperature parameters for each participant, *s*, were sampled from two group-level distributions. Participant-level learning rates *α*_*s*_ were drawn from a Beta distribution with group shape parameters *a*_1_ and *a*_2_:

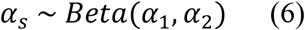

Participant-level *β*_*s*_ vectors of inverse temperature coefficients were drawn from a multivariate normal distribution with group-level means *β*_*g*_ and cross-participant covariance parameterized with standard deviations *τ* and correlation matrix Ω:

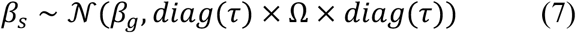

where *diag*(*τ*) is a square diagonal matrix with the elements of *τ* on the diagonal. Group-level parameters were drawn from the following hyperprior distributions to regularize estimation:

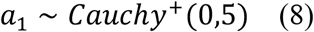

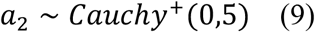

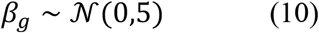

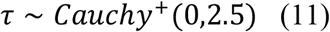

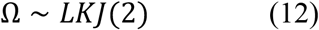

The joint posterior distribution for all parameters was sampled using Hamiltonian Markov Chain Monte Carlo (MCMC) with No-U-Turn Sampling^46^, as implemented in Stan^47^, using 4 chains with 4000 samples (2000 discarded as burn-in). Chain convergence was assessed by ensuring that all Gelman-Rubin statistics were close to 1. We compared models that included only incremental or episodic value in the choice function with those incorporating both sources of value using leave-one-out cross-validation estimated using Pareto-smoothed importance sampling^48^. The expected log pointwise predictive density (ELPD) was then computed as a measure of estimated out-of-sample predictive fit for each model. Effects are reported throughout as the mean of the posterior estimate alongside 95% credible intervals. Separate models were fit for each group (PD on medication, PD off medication, and healthy controls). Fit parameters were compared across groups using paired samples t-tests such that each patient was matched with themselves (on versus off-medication) and with independent t-tests to compare with age- and education-matched healthy controls (HC v. PD on-medication and HC v. PD off-medication). Multiple comparisons were accounted for in these matched sample analyses using the FDR correction, and adjusted q-values are reported throughout.

To analyze learning rates for each group, we initially determined the optimal learning rate for each participant’s individual sequence of outcomes using the following procedure. First, we conducted a grid search of learning rates (in units of 0.01 between 0 and 1) for each participant by i) simulating Q values for each participant, ii) computing the mean square error between the resulting sequence of Q values and whether each deck was currently lucky or not according to the task design, and iii) determining which learning rate minimized the mean square error. We then subtracted each participant’s optimal learning rate from their fit learning rate. These difference scores were then used as the outcome variable in simple linear regressions, estimated using rstanarm for each group, with an intercept as predictor in order to assess whether, at the group level, learning rates were different from optimal or not. Between group differences were then assessed using paired- (on v. off-medication) and independent- (HC v PD on-medication and HC v. PD off-medication) sample t-tests with an FDR correction for multiple comparisons.

Finally, we assessed subsequent memory performance for each group separately. Recognition memory was measured using the signal detection metric *d′*, whereas value memory was assessed as participants’ accuracy at recalling the true value of a previously seen object. Group-level effects were determined by simple linear regressions, again estimated using rstanarm, with either *d′* or value accuracy as the outcome variable and an intercept as predictor in order to assess differences from chance. *d′* was measured as the difference in z scored hit rate and false alarm rate for each participant, adjusted for extreme proportions using a log-linear rule^49^, and value memory accuracy was corrected for chance performance (accuracy - 1/6). An additional mixed effects logistic regression was used to test for an effect of the true value of previously seen objects on value memory accuracy. For this analysis, we subtracted the absolute value of 0.5 from an object’s true value and used this as the primary fixed effect alongside random slopes that varied for each participant. We chose to analyze the data in this way rather than as a quadratic effect of an object’s true value because, due to the conditional nature of this response, not all subjects had trials for every individual value, and this procedure ensured that there were adequate trials in each value bin to estimate an effect. Between group differences in *d′*, value memory accuracy, and the random slopes capturing the effect of object value on value memory accuracy for each participant were then assessed using paired- (on v. off-medication) and independent- (HC v PD on-medication and HC v. PD off-medication) sample t-tests with an FDR correction for multiple comparisons. This method was also used to assess between group differences in median reaction times as well as forward and backwards digit span scores. One patient (and their matched healthy control) was excluded from all subsequent memory analyses due to a lack of responses, and a second patient (and their matched healthy control) was excluded from only value memory analyses due to a lack of responses on these trials specifically.

For the computational model and regression analyses, fixed effects are reported in the text as the mean of each parameter’s marginal posterior distribution alongside 95% credible intervals, which indicate where 95% of the posterior density falls. Given the data and model, the probability of parameter values outside of this range is less than 0.05.

### Data availability

The data used in this study are available upon request from the corresponding author.

## Results

### Choices are best explained by a model featuring both incremental and episodic value

To assess whether participants’ responses were based on episodic memory or incremental learning, we fit a reinforcement learning model featuring separate inverse temperature parameters for model-estimated deck value and the true value of previously seen objects (see **Methods**). Additional predictors were included to capture any potential confounding biases in choice, such as a bias toward one deck color over the other, a bias toward selecting previously seen objects regardless of their value, and a perseveration bias. In this model, deck value represents personalized trial-by-trial estimates of each participant’s valuation of either deck over the course of the experiment and reflects the outcome of incremental value updating. By contrast, remembered object values were simply included as the previously experienced outcome, as participants had only one relevant past experience with an object for decisions of this type. We compared the ability of this combined reinforcement learning model to predict choices with models that included only either estimated deck value or episodic value, rather than both sources.

We found that for all groups (patients on- and off-medication and healthy controls), the combined model yielded the best estimated out-of-sample predictive performance (see **Methods** and **Supplemental Figure 1A-B**), indicating that all groups used a combination of incremental learning and episodic memory in their choices.

We next sought to examine the degree to which each group’s choices could be explained by either of these strategies, separately. We compared the performance of the two models, each of which based decisions on either incremental learning or episodic memory alone. Importantly, while episodic memory was not an available strategy on all trials, participants always had the option to base decisions on incrementally learned deck value. We therefore predicted that the incremental learning-based choice model should outperform the episodic memory-based choice model *if* participants engaged in incremental learning throughout the task. This was the case for healthy controls (expected log pointwise predictive density M=-216.723, SE=22.449; **Supplemental Figure 1C**) and for PD patients on-medication (M=-98.695, SE=20.282) but was not found for PD patients off-medication (M=-7.458, SE=17.704), indicating that behavior in the PD off group was not consistent with incremental learning. Together, these results suggest that while all groups used both sources of value to some extent, the behavior of PD patients off dopaminergic replacement medication was less consistent with an incremental learning strategy.

### Dopamine depletion decreases incremental learning but not episodic memory for choice

The model comparisons provide initial evidence that dopamine depletion in Parkinson’s Disease selectively impairs incremental learning. Next, we moved to investigate the parameters governing participants’ individual choices to further characterize this pattern. We first examined the extent to which deck decisions were well explained by estimated deck value, which provided a subject-specific predictor of incremental learning and tracked well with the reversal structure of the task (**Supplemental Figure 2**). While deck value predicted choices made by all groups to some extent (HC: β_deckvalue_ = 3.59, 95% CI = [2.258, 4.993]; PD on: β_deckvalue_ = 2.765, 95% CI = [1.318, 4.163]; PD off: β_deckvalue_ = 1.248, 95% CI = [0.559, 2.099]; **Figure 2 and 3A, Supplemental Table 2**), PD patients off-medication made less use of deck value than both healthy controls (t(25)=7.685, p<0.0001, FDR corrected) and when they were administered their dopamine replacement medication (t(25)=-9.076, p<0.0001, FDR corrected). Patients on-medication also performed similarly to healthy controls, with no significant difference in sensitivity to deck value between the on-medication and healthy control groups (t(25)=2.521, p=0.149, FDR corrected). Critically, these results indicate that dopamine depletion impaired patients’ ability to learn about reward from trial-and-error and use this information to inform their decisions. Further, they suggest that incremental learning was improved when patients were tested on their dopaminergic medication.

**Figure 2.**
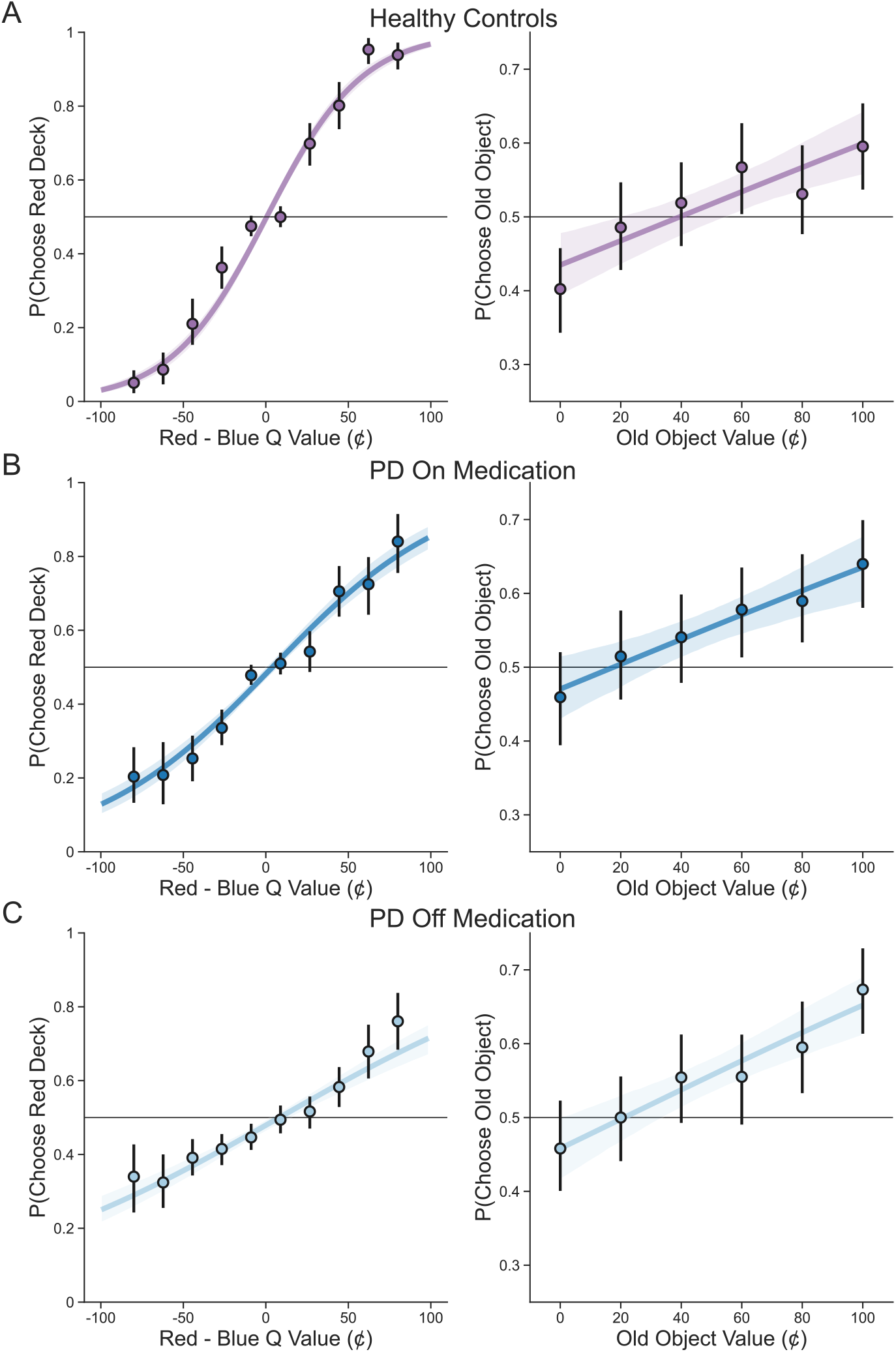
Memory-based decision-making task performance for all groups. **A)** Left: Deck learning performance for healthy controls, as indicated by the proportion of trials on which the red deck was chosen as a function of the difference between estimated value (Q) of the two decks. Right: Use of episodic value in decisions for healthy controls, as indicated by the proportion of trials featuring a previously seen (old) object on which the old object was chosen as a function of its value. The same is shown for Parkinson’s patients on (**B**) and off (**C**) medication. In all plots, each line indicates the fit of a logistic regression model to the raw data with bands representing 95% confidence intervals around this fit. Filled points represent the group average over binned data with 95% confidence intervals.

**Figure 3.**
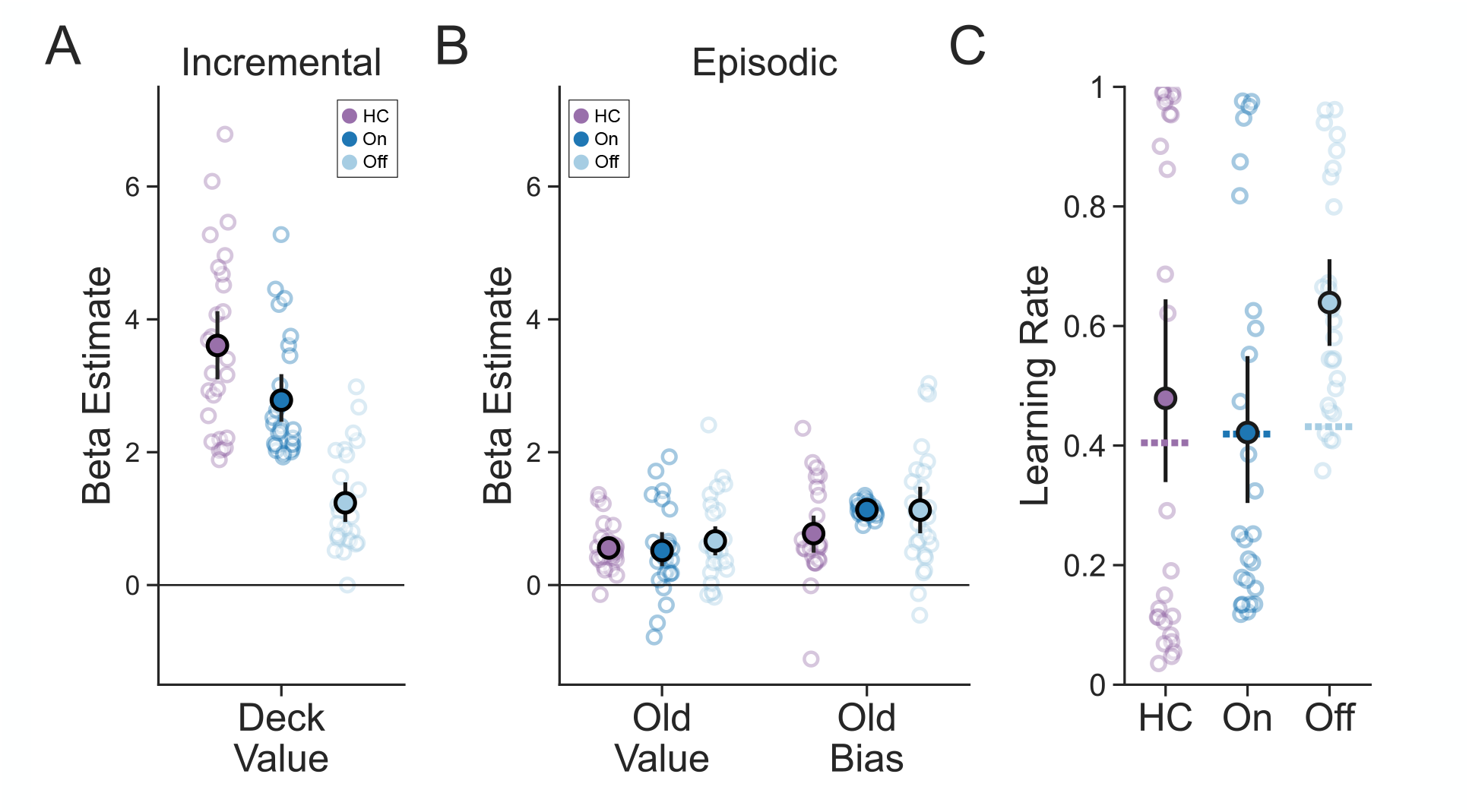
Reinforcement learning model results. To capture behavior in on the memory-based decision-making task, we fit a reinforcement learning model that incorporated both incremental and episodic value, as well as several other biases (see **Methods**), into its decisions. **A)** Inverse temperature (Beta) parameters captured sensitivity to estimated deck value. Healthy controls (in purple) and PD patients on medication (in dark blue) were more sensitive to deck value than PD patients off medication (in light blue). **B)** Further inverse temperature (Beta) parameters captured two influences on decisions related to episodic memory. The inverse temperature for old value captured sensitivity to a previously seen object’s value, whereas that for old bias captured participants’ tendency to choose previously seen objects regardless of their value, which was a form of recognition memory bias. Filled points represent group-level mean parameter estimates with 95% confidence intervals and empty points represent individual parameter estimates. **C)** Learning rate estimates for each group from the combined reinforcement learning model. PD patients off medication had higher learning rates than patients on medication, but not healthy controls. Further, PD patients off medication were the only group with a learning rate that was different from the optimal learning rate (dotted lines), and the extent of this suboptimality was again greater than for patients on medication, but not healthy controls. Filled points represent group level average learning rate estimates, empty points represent individual subject estimates, and error bars represent 95% confidence intervals.

We next asked whether dopamine depletion had any impact on patients’ use of episodic memory. Specifically, if all patients were performing well, and the off-medication patients were not as effective at using an incremental learning strategy, were they instead relying on episodic memory? To answer this question, we examined whether decisions on trials featuring a previously seen object were well explained by that object’s value, which provided a measure of episodic memory use throughout the task. Object value predicted choices made by all groups (HC: β_oldvalue_ = 0.559, 95% CI = [0.139, 1.027]; PD on: β_oldvalue_ = 0.516, 95% CI = [0.017, 1.043]; PD off: β_oldvalue_ = 0.657, 95% CI = [0.149, 1.191]; **Figure 2 and 3B**), and there were no differences between patients on or off their medication and healthy controls (HC v. PD off: t(25)=-0.733, p=1.0, FDR corrected; HC v. PD on: t(25)=0.266, p=1.0, FDR corrected; PD on v. PD off: t(25)=0.953, p=1.0, FDR corrected). In addition to sensitivity to object value, we also observed that participants’ choices tended to be biased by previously seen objects regardless of their value. This bias was similarly predictive of choice across all groups (HC: β_oldbias_ = 0.773, 95% CI = [0.348, 1.214]; PD on: β_oldbias_ = 1.133, 95% CI = [0.87, 1.396]; PD off: β_oldbias_ = 1.121, 95% CI = [0.622, 1.646]) and there was no significant difference between patients on or off their medication and healthy controls (HC v PD off: t(25)=-1.561, p=0.776, FDR corrected; HC v. PD on: t(25)=-2.560, p=0.149, FDR corrected; PD on v. PD off: t(25)=-0.041, p=1.0, FDR corrected). Together, these results indicate that while PD patients off-medication engaged in less incremental learning, their use of episodic memory remained unaltered. Further, administering dopamine replacement medication had very little impact on episodic memory usage.

### Dopamine remediates a suboptimal rate of incremental learning in PD patients

Having established that dopamine depletion selectively impairs incremental learning in our task, we next moved to better characterize the nature of this deficit by examining the learning rate parameter of our combined reinforcement learning model. In incremental learning models of this type, which employ a Rescorla-Wagner update, the learning rate is responsible for controlling the magnitude to which reward prediction errors on each trial override the previously stored value for each eligible cue. Higher learning rates effectively amount to larger exponential discounting of past rewards. Critically, the optimal learning rate for a task is dependent on that task’s volatility— when change is more frequent, a higher learning rate is more optimal^50–52^. Achieving an optimal learning rate is possible, and desirable, only if reward prediction errors remain intact, as this is the signal over which the learning rate operates. We therefore hypothesized that if dopamine depletion impairs reward prediction error signaling in PD patients off-medication, then their learning rate should be suboptimal relative to patients on-medication.

In support of this idea, patients tested off- vs. on-dopaminergic medication significantly differed in their learning rates (t(25)=3.533, p=0.009, FDR corrected; **Figure 3C, Supplemental Table 2**). To gain more insight into the nature of this difference, we next examined whether each group’s learning rates differed from the learning rate that was optimal in the present task. When tested off-medication, patients updated faster than was optimal (M = 0.207, 95% CI = [0.120, 0.294]). When tested on-medication, however, patients learned at an optimal rate (M = 0.002, 95% CI = [-0.136, 0.135]), and did so significantly more than when they were tested off-medication (t(25)=3.131, p=0.024, FDR corrected). This result suggests that restoring reward prediction error signaling allowed patients to set their learning rate to a level that was more optimal. Separately, while at the group-level healthy controls also learned at an optimal rate (M = 0.074, 95% CI = [-0.098, 0.241]), there was a large degree of variability across individuals and their overall learning rates were not different from patients off- (t(25)=-1.776, p=0.225, FDR corrected) or on-medication (t(25)=0.556, p=1.0, FDR corrected). Together, these results suggest that achieving an optimal rate of incremental learning is aided by the presence of phasic dopamine.

### Dopamine enhances subsequent memory for extreme values with little effect on recognition memory

Following the memory-based decision-making task, we immediately tested participants’ ability to recall a subset of the trial-unique objects (**Figure 1A**). Subsequent memory was assessed in two ways. First, we asked participants to identify whether or not they recognized objects from the prior task. Second, if they indicated that they both recognized and had chosen an object, we asked participants whether they remembered its value. This task provided us with an opportunity to further probe any potential differences in episodic memory formation due to the presence or absence of dopamine in PD patients.

All groups recognized previously seen objects well above chance (HC: β_intercept_ = 1.400, 95% CI = [1.061, 1.749]; PD on: β_intercept_ = 1.187, 95% CI = [0.915, 1.462]; PD off: β_intercept_ = 1.377; 95% CI = [1.017, 1.739]; **Figure 4**). There were also no differences between the groups (HC v. PD off: t(24)=0.072, p=1.0, FDR corrected; HC v. PD on: t(24)=0.982, p=0.910, FDR corrected; PD on v. off: t(24)=1.317, p=0.910, FDR corrected). These results suggest that dopamine played little-to-no role in recognition memory for individual object stimuli.

**Figure 4.**
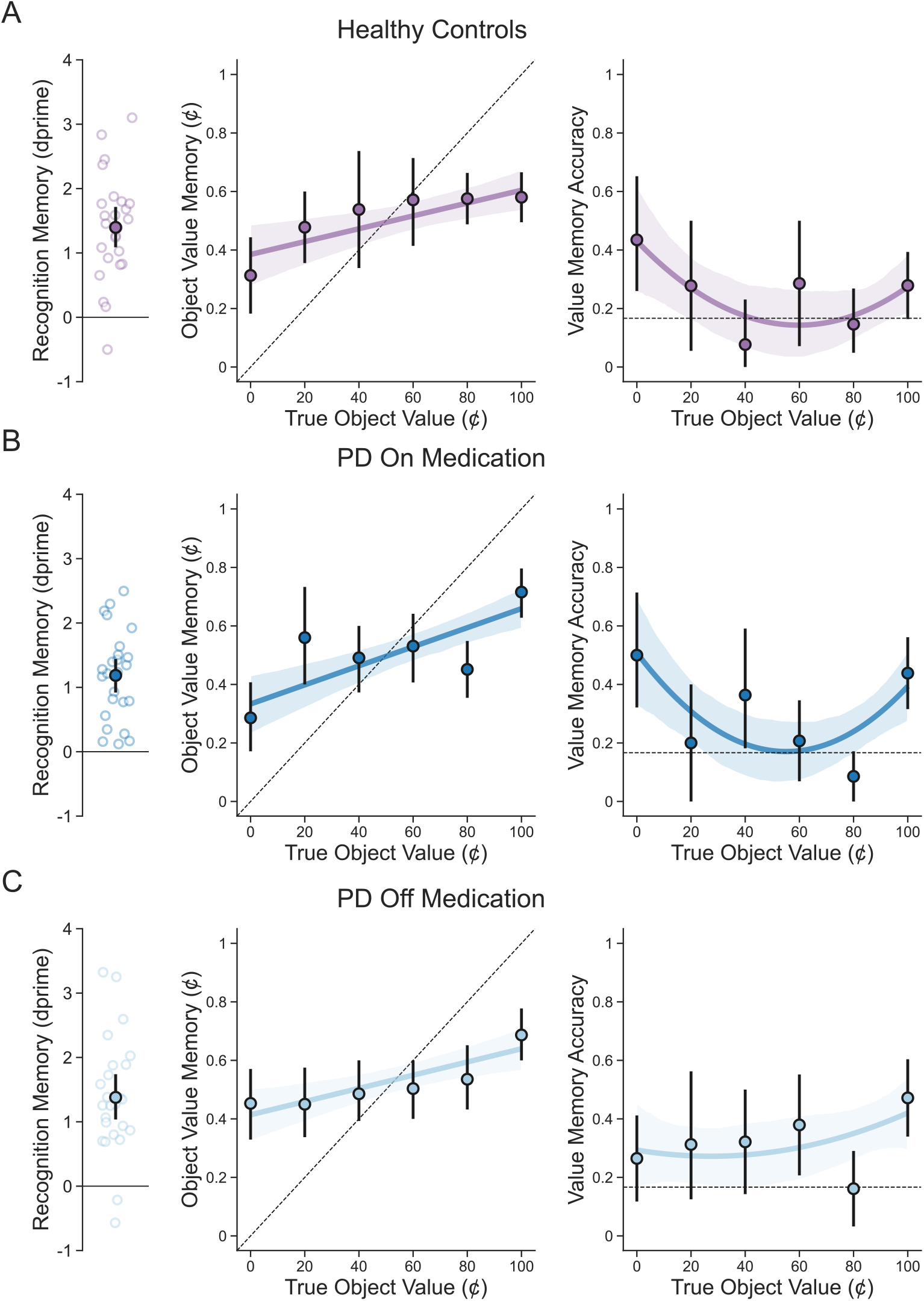
Subsequent memory task performance for all groups. **A)** Left: Recognition memory performance on the subsequent memory task as captured by the signal detection metric dprime for healthy controls. Middle: Value memory performance visualized here as the relationship between healthy controls’ memory for object value and an object’s true value. Right: Value memory performance visualized the relationship between accuracy (whether the correct value was remembered or not) and an object’s true value. The same is shown for Parkinson’s patients on (**B**) and off (**C**) medication. For the left panel, filled points represent the group average with a 95% confidence interval and empty points represent individual participant means. For the middle and right panels, filled points represent the group average over binned data with 95% confidence intervals and each solid line with bands indicates the fit of regression models to the raw data with bands representing 95% confidence intervals around this fit.

We further probed participants’ ability to remember the value associated with objects that were both recognized and chosen. Because this requires a more discerning response about associations and actions, we reasoned that accurate value memory was likely indicative of more episodic-like subsequent memory than the test of recognition memory alone. We first assessed whether participants were able to recall the value of chosen objects above chance. While patients both on-(β_intercept_ = 0.156, 95% CI = [0.047, 0.264]; **Figure 4**) and off-medication (β_intercept_ = 0.153, 95% CI = [0.067, 0.237]), were indeed able to recall object value, controls were no different from chance (β_intercept_ = 0.092, 95% CI = [-0.011, 0.196]). While one interpretation of this result was that healthy controls may have a lower quality of memory for value overall, there were no differences between healthy controls and patients when we compared between groups (HC v PD off: t(23)=-0.971, p=1.0, FDR corrected; HC v PD on: t(23)=-0.907, p=1.0, FDR corrected; PD on v PD off: t(23)=-0.046, p=1.0, FDR corrected).

We next asked whether the value of each object mattered for memory, given that dopamine is thought to prioritize memories associated with motivationally significant events^36,53–55^. Here we reasoned that the value of objects that were more motivationally relevant for choice (i.e., those with more extreme value) would be better remembered, and that the presence or absence of dopamine would modulate this effect. We indeed found that PD patients tested on dopamine replacement had greater recall of object value for objects with more extreme value (β_value_ = 0.548, 95% CI = [0.069, 1.118]), and that this effect was absent when patients were off-medication (β_value_ = 0.111, 95% CI = [-0.195, 0.424]; PD on v. PD off: t(23)=-5.486, p<0.0001, FDR corrected).

While healthy controls also did not demonstrate this modulation of value memory by motivational significance at the group-level (β_value_ = 0.393, 95% CI = [-0.138, 0.952]), there was no difference in this effect when healthy controls were compared to PD patients on-medication (t(23)=-1.664, p=0.110, FDR corrected). Healthy controls also had better value memory relative to PD patients-off medication (t(23)=14.121, p<0.0001, FDR corrected). Overall, these results suggest that while the presence or absence of dopamine had little effect on episodic memory capacity when tested with either a simple recognition memory test or a test for associated value memories, dopamine leads to the selective prioritization of motivationally relevant value information in memory.

### Controlling for effects of motor deficits and other cognitive and psychiatric variables

To confirm that differences reported here for PD patients were not due to general motor impairment caused by PD, we also assessed the relationship between motor symptoms, as measured by UPDRS, and sensitivity to incrementally learned value. There was no correlation between the UPDRS scores of PD patients and task performance (**Supplemental Table 1)**. To better isolate effects of improvement due to dopamine, we also examined the relationship between each patient’s difference in UPDRS scores (on – off medication) and incremental value sensitivity (on – off medication), and found no correlation (r=0.201; p=0.325). Lastly, PD patients off-medication and healthy controls demonstrated no differences in reaction time on the memory-based decision-making task (t(25)=-0.995, p=1.0, FDR corrected), suggesting that any behavioral differences reported here cannot be attributed to slower motoric responses in PD patients.

Finally, all participants completed a battery of cognitive and psychiatric assessments including the MOCA, forwards and backwards digit span tasks, and the GDS and Starkstein apathy scale (see **Methods**). All participants had a MOCA score >=26 indicating no cognitive impairment and there was no difference between HC and PD patients in MOCA scores or digit span performance (**Table 1**). It is therefore unlikely that the differences observed in the present study were due to overall differences in cognitive or working memory impairment. While PD patients had greater apathy scores than healthy controls, as has been reported previously^56^ (**Table 1**), apathy was not related to incremental learning sensitivity. Furthermore, none of the cognitive, psychiatric, or demographic variables we collected were significantly correlated with any of the learning and memory measures in the task (**Supplemental Table 1**).

## Discussion

The majority of studies on reward-based learning have focused on how people update an option’s value gradually over time. However, the extent to which learning and decisions are based instead on single past encounters has been a primary focus of much recent work.^9–11,23,24,26^ While computational theory has suggested that individual events may be particularly useful when experience with a task is limited,^29,57^ experimental studies have demonstrated that they are relied upon even throughout tasks that can be solved normatively with incremental learning alone.^9–11^ This suggests that information encoded in a single shot with episodic memory is commonly recruited for reward-based learning and decision-making. One prevailing hypothesis is that episodic memory is needed specifically for decisions based on individual events rather than those based on incrementally constructed summaries over many events. However, this dissociation has yet to be shown empirically. Studying PD patients, who are known to be impaired at learning from feedback due to loss of striatal dopamine,^5,58,59^ provides an avenue for investigating this dissociation. Using a task that separately measured the contributions of episodic memory and incremental learning to decisions, we found that PD patients were able to base their choices on single experiences but were impaired at choosing based on incremental learning. Furthermore, PD patients’ subsequent episodic memory for individual events remained intact following decision-making. Our findings therefore suggest that reward-based learning, when it relies upon episodic memory, can occur in the absence of dopamine-replacement medication in PD. They further demonstrate that when contributions from episodic memory are properly controlled for, PD patients remain impaired at reward learning from trial-and-error.

The medial temporal lobe (MTL), and particularly the hippocampus, may instead support reward-based learning that relies upon episodic memory. Indeed, while countless studies have demonstrated that the MTL is unequivocally necessary for episodic memory,^60,61^ it has also been shown that the MTL is recruited for decisions to which episodic memory may contribute.^21,22,62^ There is further evidence that the MTL plays an important role in both value-based decision-making and learning from feedback. First, patients with MTL damage show less consistent value-based choices when compared to healthy controls,^21,22^ and it has been demonstrated that BOLD activity in the hippocampus increases when deliberation between choice options takes more time.^21^ Second, neurons in the hippocampus code for rewarding outcomes throughout learning.^63–65^ In humans, evidence from both fMRI^66^ and lesion^67^ studies has shown that the MTL is needed specifically when feedback is delayed and must be bound across time. While these studies all reflect the sorts of relational computations thought to be performed by the MTL^68,69^ and underlying episodic memory^70,71^, recruiting episodic memory is not explicitly necessary in any of this work. In fact, to the best of our knowledge, little work has been done to test whether the MTL is required for decisions in which intact episodic memory is unambiguously required for adequate performance. This is, in part, due a lack of tasks which can cleanly separate the contributions of episodic memory from other memory systems. One avenue for future work would be to test for a causal role of the MTL in value-based decisions that are clearly based on episodic memory using the present task in patients with MTL damage.

It has been suggested that episodic memory may in some cases be recruited more in order to compensate for deficits in incremental learning. We did not find support for this in the present work. This hypothesis relates to a more general idea in the field of cognitive control: that the brain may judiciously adopt different decision strategies under the circumstances for which each is likely to produce the most rewarding choices. Most notably, this logic has been used to explain how the brain arbitrates between deliberative and habitual decisions,^72,73^ and was recently extended to investigate competition between incremental learning and episodic memory using a variation of the task presented here.^24^ In that study, it was found that decisions were indeed guided by episodic memory more often when incremental learning yielded unreliable estimates of value. In the present work, we found that PD patients off-dopamine replacement medication were selectively impaired at incremental learning, and that they recruited episodic memory for decisions to the same extent as healthy controls.

Given that impaired incremental learning should yield particularly unreliable estimates, this finding is somewhat surprising. There are at least two possible explanations. First, Nicholas et al., (2022) found that tracking uncertainty around incrementally constructed estimates was important for successful arbitration with episodic memory. Because this type of summary statistic is fundamentally built upon reward prediction error signaling, it is reasonable to assume that patients off-medication were not able to compute it. Second, in our task, an ideal observer with perfect episodic memory would always benefit from using it to make decisions. Yet we find that participants are far from this ideal: they are biased by recognition and limited in their ability to remember object identity and value. The fact that all groups had a similar capacity for remembering individual objects in our subsequent memory test suggests that they may be performing at, or near, the limits of episodic memory. The presence of such a ceiling effect may therefore make it difficult to adequately measure any compensatory usage of episodic memory in the present task.

Relatedly, while initially it was thought that PD patients demonstrate few issues with episodic memory,^4^ it has since emerged that PD patients, even in early stages, are often impaired in this domain.^74^ Specifically, PD patients have been reported to show large deficits on tests on both immediate and delayed measures of free recall,^75^ and a number of studies have found that recognition memory is similarly affected,^12–16^ although evidence is mixed on this point.^33–36^ While the exact reasons for these differences are not currently known, recent evidence suggests that atrophy in the MTL is more common with PD than previously thought,^74,76^ and that MTL atrophy is related to memory impairments in PD.^76,77^ Although episodic memory deficits and MTL atrophy are more common with later disease progression, both have been reported during earlier stages of PD.^74^ Here we find no evidence for episodic memory impairment in PD patients, both on and off their dopamine replacement medication, relative to healthy controls. Not only were patients able to base choices on their memory for single experiences, but they were also able to both recognize and remember the value of these individual cues when tested following a short delay. These findings therefore provide further support for intact episodic memory during the mild-to-moderate stages of PD.

Furthermore, we found that administering dopamine replacement medication to PD patients improved their memory for the value of individual objects when it was most relevant for choice. This result joins recent work in PD patients^36^ to suggest that dopamine aids in prioritizing the formation of long-term memories that are associated with motivationally significant.^53–55^ While one past study investigated the effects of dopamine on incidental information presented alongside reward,^36^ here we extend these findings by demonstrating that dopamine strengthens subsequent memory for the motivational information associated with single events regardless of its valence. Critically, in the present experiment, all individual cues and their value were relevant for guiding choice, which allowed us to examine dopamine’s effects on memory for information that was not merely incidental. Further, as in Sharpe et al. (2020), we also found relatively no effect on the overall capacity of both recognition and value memory, regardless of patients’ medication status. These findings therefore lend additional support to the idea that dopamine effects what, as opposed to how much, is stored in memory.

Another consideration related to the effect of PD on incremental learning that we found here is that the exact nature of this impairment is currently not well understood. While studies have typically assumed that the absence of striatal dopamine in PD patients leads to deficiencies in incremental learning, recent findings^32^ have suggested that it may instead impair learning that is more goal-directed and reliant on building models of task structure. Work in rodents has similarly demonstrated that phasic dopamine signals are important for learning associations between stimuli to support model-based inference.^78,79^ In light of these findings, one possibility is that the types of paradigms typically used to measure incremental learning may instead measure model-based learning, thereby masking the true source of PD patients’ impairment. Two potential avenues for this misattribution may be that participants could develop rules and/or use working memory to guide their choices.^32,80–82^ Although the way that incremental learning was operationalized in our experiment was, by itself, no different from a typical two-armed bandit task often seen in the literature, there is evidence that neither of these potentially model-based approaches were used. First, we found no relationship between patients’ sensitivity to learned deck value and our measures of working memory. Second, past work^24^ deploying a variant of the same task in healthy adults found that participants were more likely to engage in incremental learning than rule-based inference. Regardless, while here we have shown that PD patients are capable of using individual trials to guide their choices, suggesting that this strategy does not contribute to their inability to learn from trial-and-error, future work will be required to similarly disentangle contributions of model-based learning as well.

We found that PD patients’ incremental learning impairment manifested as both decreased sensitivity to learned deck value and also a suboptimal rate of learning. Importantly, an altered learning rate in the absence of phasic dopamine is expected if this signal is responsible for encoding reward prediction error. This is because, in standard models of incremental learning like the type we employed here, cue values are updated by the product of the learning rate and the reward prediction error.^20^ While past work has found that PD patients off-medication may update value either more^83^ or less^84^ quickly than when on-medication, we note here that the optimal learning rate differs from task-to-task. In particular, what learning rate will be best is modulated by a task’s baseline level of volatility, or how frequently changes in the mapping between actions and outcomes occur.^50–52^ While we found that PD patients off-medication updated more rapidly than when on-medication, critically, when patients were administered their dopamine replacement medication they learned more optimally in response to the statistics of our particular task environment. This result provides evidence that optimal learning is more likely in the presence of intact reward prediction error signaling.

In conclusion, these results demonstrate that striatal dopamine depletion in PD impairs incremental learning from trial-and-error with no effect on the recruitment of episodic memory for choice. Dopamine replacement remediated this deficit while enhancing subsequent memory for the value of motivationally relevant stimuli. These effects suggest both that decisions based on the value of single experiences can be made in the absence of striatal dopamine, and that PD patients remain impaired at learning about reward from trial-and-error when episodic memory is properly controlled for. By probing the role of these separate memory systems in a single decision-making task, these findings shed light on the extent to which either are modulated by the presence or absence of dopamine in Parkinson’s disease.

## Abbreviations

PD: Parkinson’s disease
HC: Healthy control
MoCA: Montreal cognitive assessment
GDS: Geriatric depression scale
UPDRS: Unified Parkinson’s disease rating scale

## Supplemental Material

**Supplemental Figure 1.**
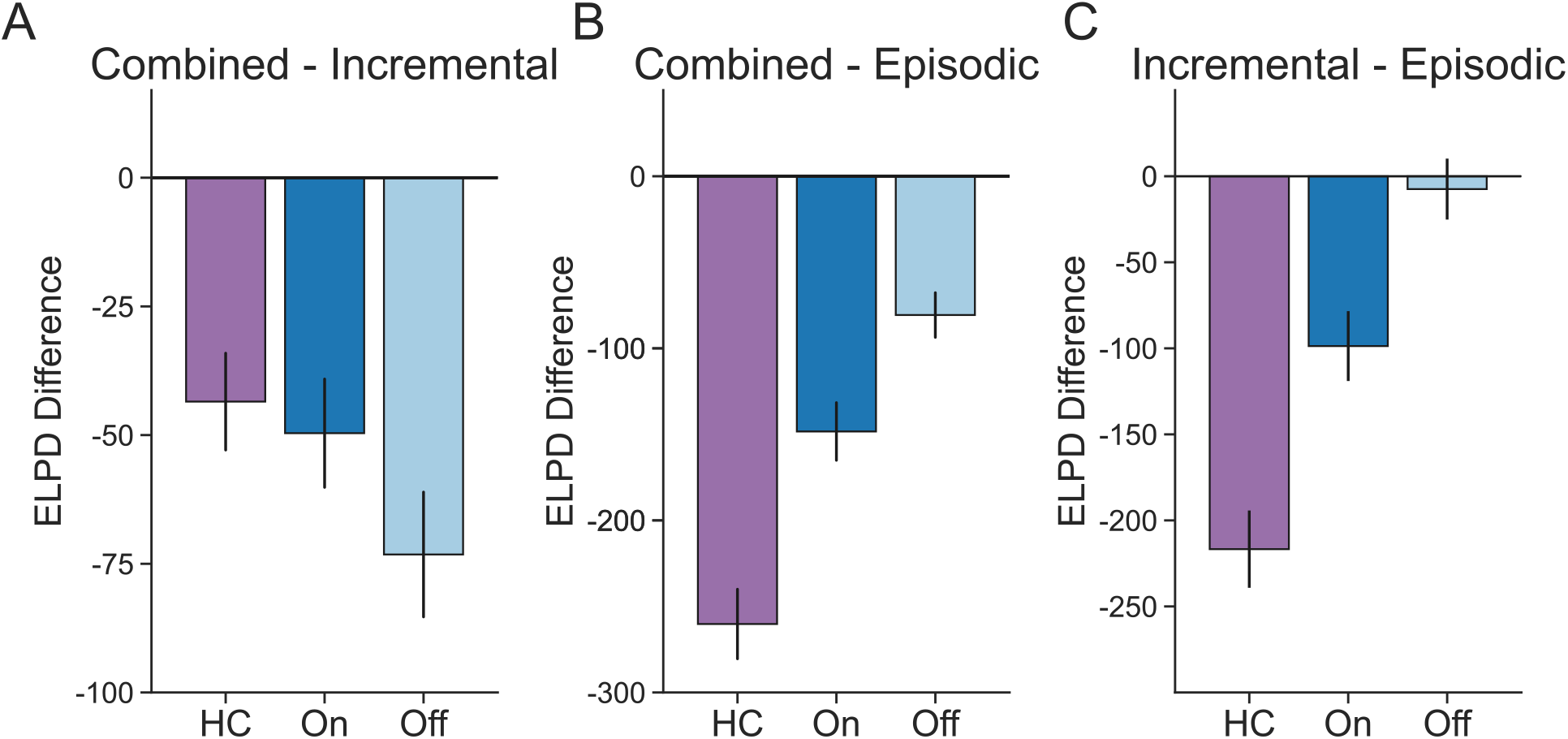
Full model comparison. The combined reinforcement learning model performed best for all groups, as measured by having the highest expected log pointwise predictive density (ELPD) compared to models that included only incremental (shown in **A**) or episodic (shown in **B**) influences on choice. ELPD is a measure of estimated out-of-sample fit, and more negative numbers in the difference between two models indicate a better fit for the first model. **C)** While the combined reinforcement learning model best explained choices in all groups, we also compared the performance of models that incorporated either only sensitivity to incremental value or sensitivity to episodic value to capture the extent to which choices could be explained by either. Choices made by healthy controls and PD patients on medication were, in general, better explained by incremental learning than by episodic memory, but this was not the case for PD patients off medication.

**Supplemental Figure 2.**
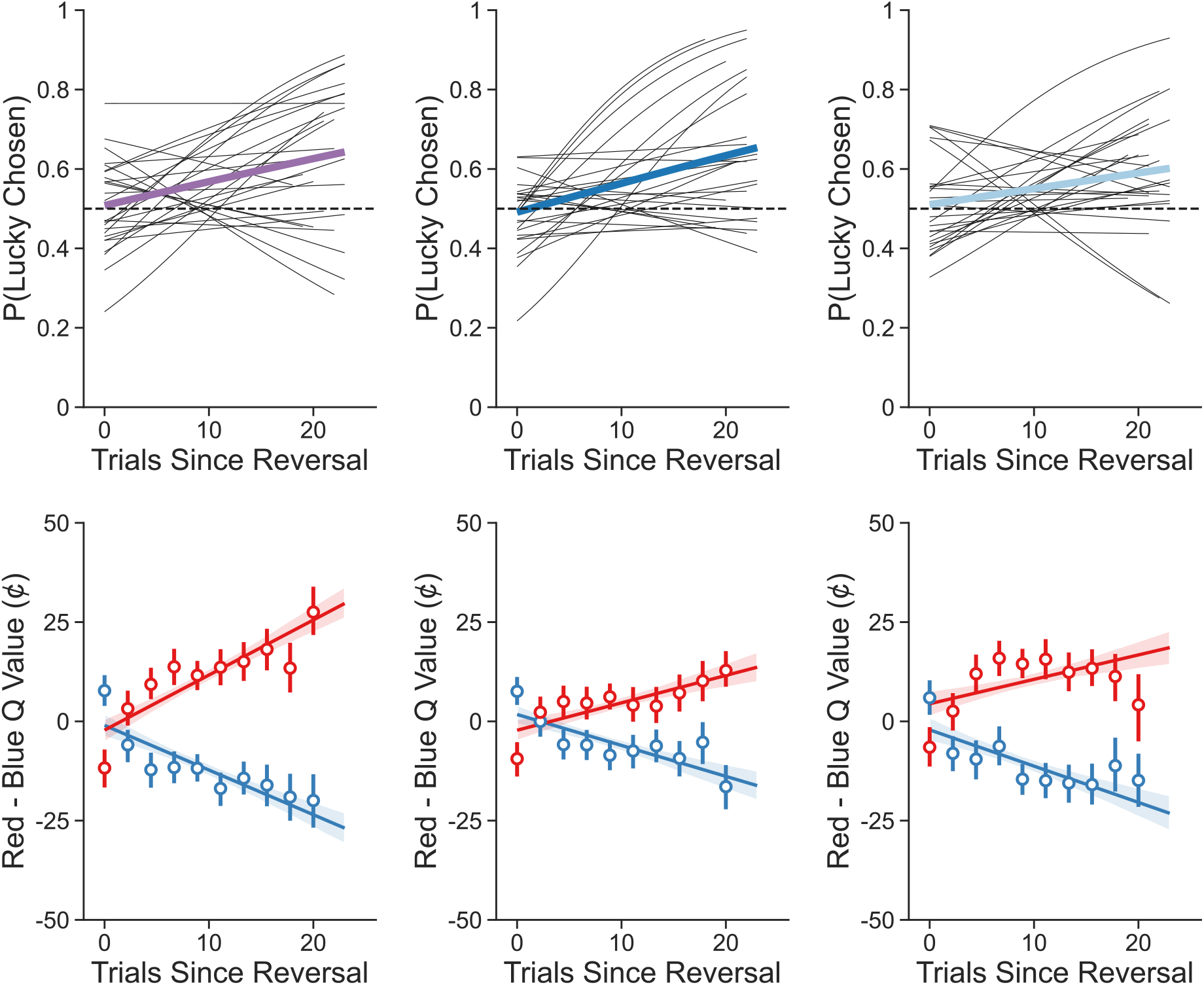
Deck learning behavior as a function of the number of trials since a reversal in deck value. **Top**: Participants tended to choose the currently lucky deck more often with more experience following a reversal. Group-level estimates of a linear regression model are shown in bold (Healthy controls: purple, PD patients on medication: dark blue, PD patients off medication: light blue) and individual subject estimates are shown as thin lines. **Bottom**: Estimated deck value from the combined reinforcement learning model tends to track with trials since reversal in all groups both when the red deck is lucky (shown in red) and when the blue deck is lucky (shown in blue). Linear regression fits are shown as lines, binned data with group averages shown as points, and error bars and bands (where shown) represent 95% confidence intervals.

**Supplemental Table 1.**
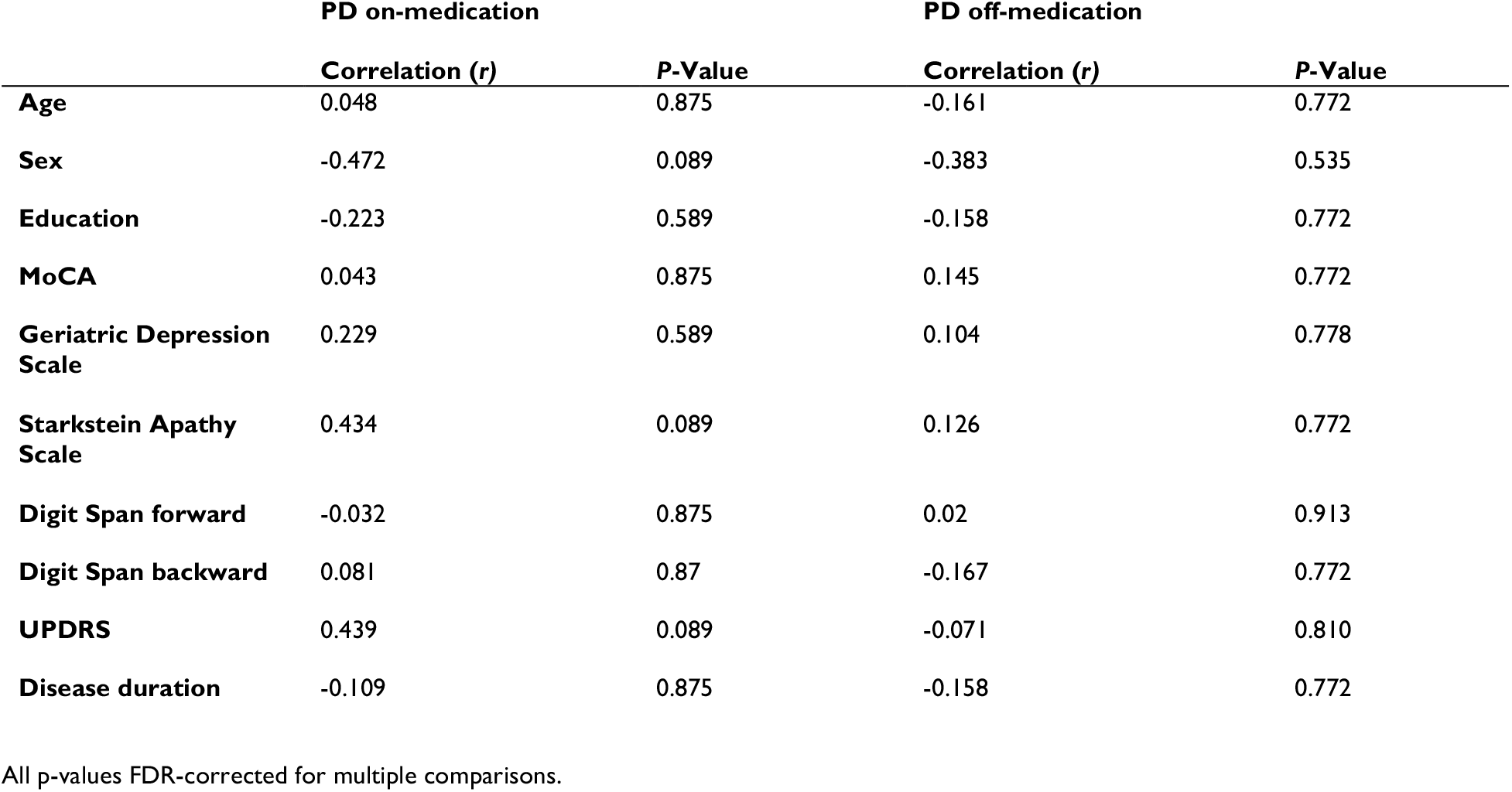
Correlations between PD participant-level measures and incremental value sensitivity (as estimated by the combined choice reinforcement learning model)

|  | PD on-medication |  | PD off-medication |  |
| --- | --- | --- | --- | --- |
|  | Correlation (*r*) | P-Value | Correlation (*r*) | P-Value |
| **Age** | 0.048 | 0.875 | -0.161 | 0.772 |
| **Sex** | -0.472 | 0.089 | -0.383 | 0.535 |
| **Education** | -0.223 | 0.589 | -0.158 | 0.772 |
| **MoCA** | 0.043 | 0.875 | 0.145 | 0.772 |
| **Geriatric Depression Scale** | 0.229 | 0.589 | 0.104 | 0.778 |
| **Starkstein Apathy Scale** | 0.434 | 0.089 | 0.126 | 0.772 |
| **Digit Span forward** | -0.032 | 0.875 | 0.02 | 0.913 |
| **Digit Span backward** | 0.081 | 0.87 | -0.167 | 0.772 |
| **UPDRS** | 0.439 | 0.089 | -0.071 | 0.810 |
| **Disease duration** | -0.109 | 0.875 | -0.158 | 0.772 |

All p-values FDR-corrected for multiple comparisons.

**Supplemental Table 2.**
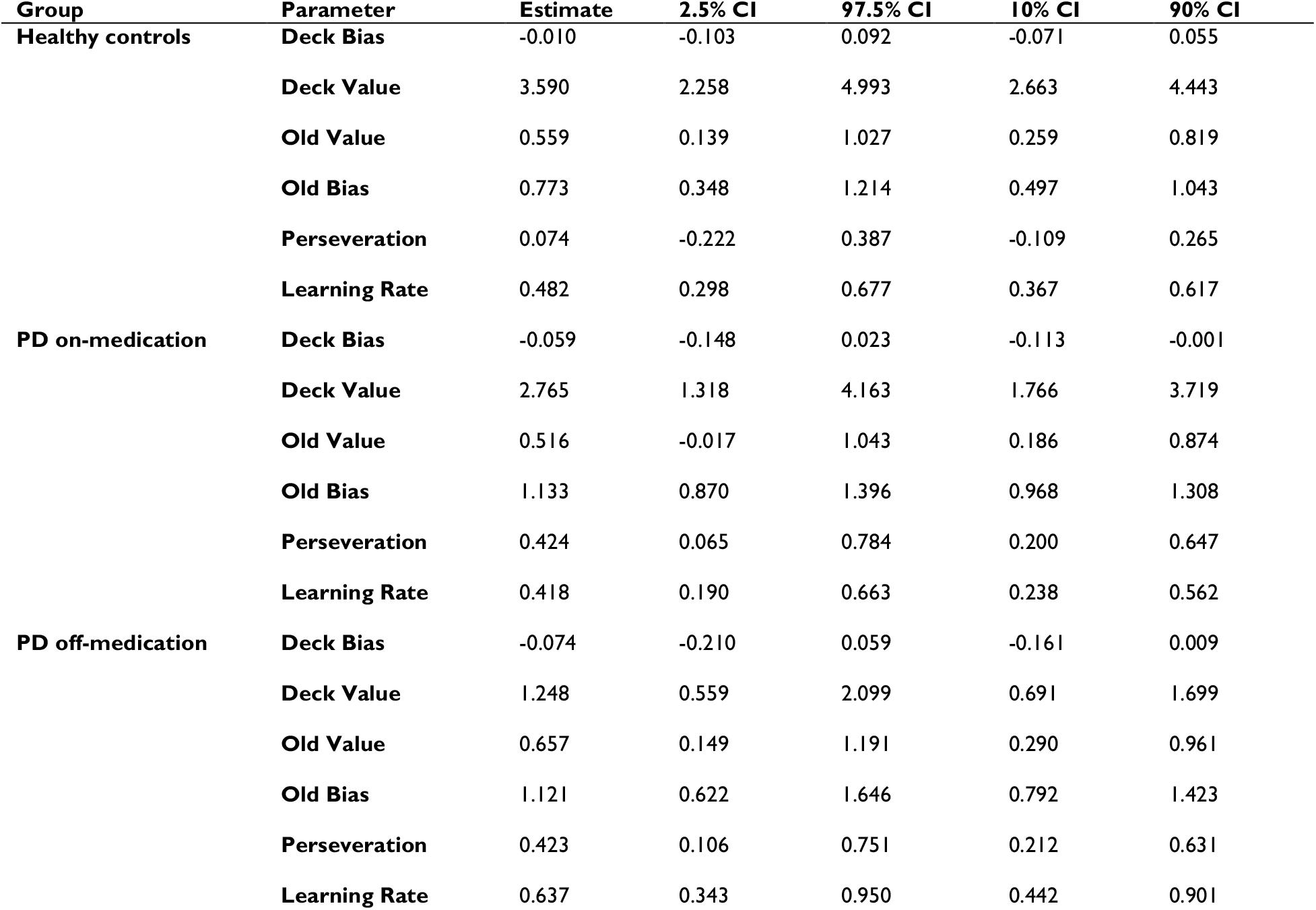
Group-level parameter fits from the combined reinforcement learning model. Parameters are reported as the mean of the posterior distribution alongside 95% and 80% credible intervals.

